# Facetto: Combining Unsupervised and Supervised Learning for Hierarchical Phenotype Analysis in Multi-Channel Image Data

**DOI:** 10.1101/722918

**Authors:** Robert Krueger, Johanna Beyer, Won-Dong Jang, Nam Wook Kim, Artem Sokolov, Peter K. Sorger, Hanspeter Pfister

## Abstract

Facetto is a scalable visual analytics application that is used to discover single-cell phenotypes in high-dimensional multi-channel microscopy images of human tumors and tissues. Such images represent the cutting edge of digital histology and promise to revolutionize how diseases such as cancer are studied, diagnosed, and treated. Highly multiplexed tissue images are complex, comprising 10^9^ or more pixels, 60-plus channels, and millions of individual cells. This makes manual analysis challenging and error-prone. Existing automated approaches are also inadequate, in large part, because they are unable to effectively exploit the deep knowledge of human tissue biology available to anatomic pathologists. To overcome these challenges, Facetto enables a semi-automated analysis of cell types and states. It integrates unsupervised and supervised learning into the image and feature exploration process and offers tools for analytical provenance. Experts can cluster the data to discover new types of cancer and immune cells and use clustering results to train a convolutional neural network that classifies new cells accordingly. Likewise, the output of classifiers can be clustered to discover aggregate patterns and phenotype subsets. We also introduce a new hierarchical approach to keep track of analysis steps and data subsets created by users; this assists in the identification of cell types. Users can build phenotype trees and interact with the resulting hierarchical structures of both high-dimensional feature and image spaces. We report on use-cases in which domain scientists explore various large-scale fluorescence imaging datasets. We demonstrate how Facetto assists users in steering the clustering and classification process, inspecting analysis results, and gaining new scientific insights into cancer biology.

## 1 Introduction

A great number of diseases, including virtually all cases of human cancer, are diagnosed by histological analysis of tissue samples, most commonly by anatomic pathologists. These samples are acquired by biopsy or surgical resection and subsequently fixed, sectioned and stained prior to examination by bright-field microscopy. Histology plays a bigger role in the diagnosis and treatment of cancer than DNA sequencing or genomic analysis. Despite this, contemporary digital technologies have not yet had much impact on anatomic pathology. There is rapidly growing interest in bio-medicine in the use of image recognition algorithms, particularly those based on deep learning, to improve the quality and reduce the cost of diagnosis [29]. This is particularly true in the case of digital histology, which is being transformed by the introduction of multiplexed imaging methods that can provide precise molecular information on cells and their constituents. For example, the Cyclic Immunofluorescence (CyCIF) approach used in this work [39, 40], generates up to 60-plex (60 channel) images of tumors and tissues at sub-cellular resolution. In such an image, each channel is acquired by staining a tissue section with antibodies that detect specific protein antigens. This makes it possible to measure the levels and locations of individual proteins and their modified forms. Information on protein levels and localization are diagnostic of cell type (i.e., cancer, immune or supporting stroma) and state (i.e., dividing, quiescent or dying). Multiplexed tissue imaging promises to combine the proven power of histology with the molecular detail hitherto provided by genomics, which currently lacks spatial resolution, in determining whether a tumor is aggressive or benign and whether it is responding to therapeutic drugs. Analyzing the data generated by multiplexed tissue imaging is the primary challenge facing widespread adoption of the method. CyCIF datasets can contain 60 or more image channels, each 30k × 30k pixels in size, with up to 10^6^ or more cells of dozens of different types. Much of the knowledge needed to interpret such images is exclusively in the brains of anatomic pathologists, who are extraordinarily skilled at identifying the hallmarks of disease by eye. These factors make data analysis very challenging. It requires joint analysis of images and image-derived data and means to tap the deep knowledge of pathologists. Few software tools currently exist to assist in this task, and they rarely support a combined visual and computational analysis.

In this paper, we describe Facetto, a new software tool for interactive phenotypic analysis of large and high-dimensional image data. The overall goal of Facetto is to provide an environment in which human experts can efficiently interact with images and image-derived data and train algorithms to perform most routine image processing tasks. This will allow a much wider range of individuals to interpret multiplex images and free pathologists to review the most salient findings to advance cancer research and improve differential diagnosis. Facetto was developed using an iterative user-centered design process with pathologists, oncologists, and computational biologists as collaborators. Facetto supports any type of high-dimensional image data including those compatible with the Bioformats Standard [8] and the wide variety of emerging tissue imaging methods [9], but this paper concentrates on multi-channel fluorescent images in the OME-TIFF format [2] acquired by CyCIF. The software integrates expectation maximization (EM) clustering and a convolutional neural network (CNN) into an interactive image exploration interface. In a typical iterative analysis process, users leverage clustering to discover and isolate new cell types and then feed the results of clustering to train classifiers which are then used to assign labels to new image data. The visual exploration of the data is supported by several different rendering modes within an image viewer and by multiple linked feature space visualizations (see Fig. 1).

**Fig. 1.**
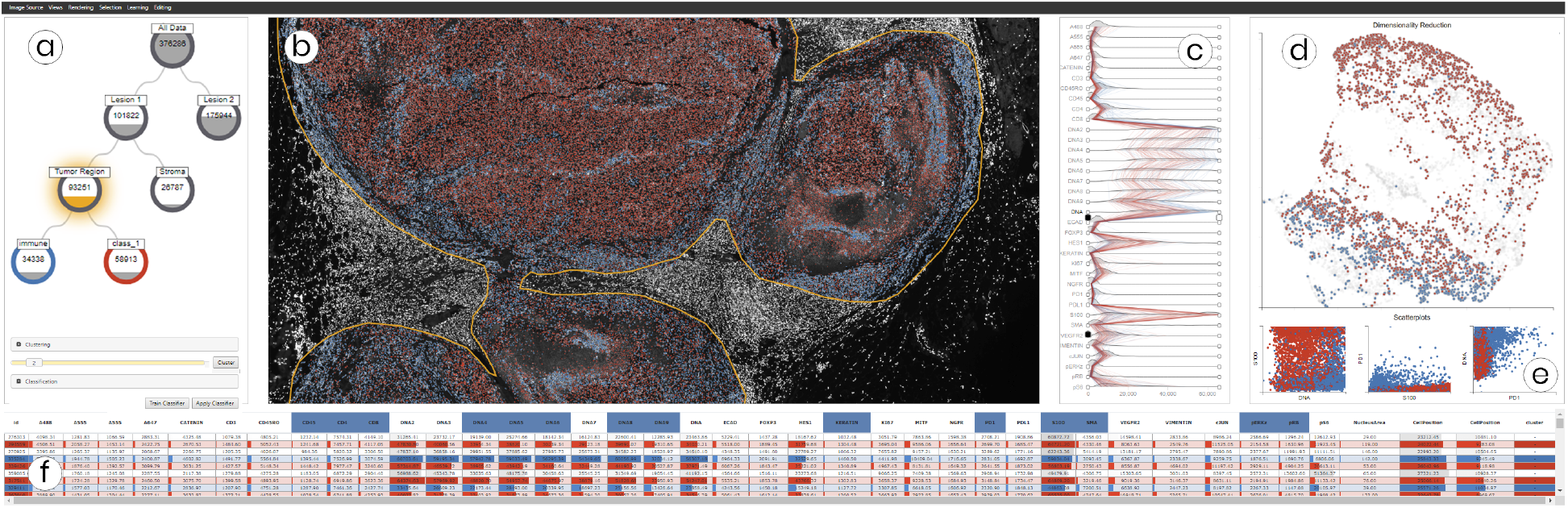
Multiple coordinated views in Facetto for interactive and hierarchical phenotype analysis of 36-channel image data (image resolution: 31,616 × 22,272 pixels; raw image size: 49.8 GB). (a) Phenotype tree resulting from hierarchical data filtering and cell calling. (b) Multi-channel visualization of high-resolution CyCIF image data showing the current clustering and classification results. (c) Ridgeplot of high-dimensional feature data to steer visual analysis and data filtering. (d) UMAP projection of the sampled feature space of cells, colored by cluster ID. (e) Scatterplots showing feature value correlations. (f) Table view of all cells and their features.

The current work makes three primary contributions. First, it applies unsupervised and supervised learning in a novel way to the task of identifying cell types and subtypes. Second, it uses a set-based data-tree visualization to hierarchically subdivide (facet) large image and feature datasets into smaller subsets and to annotate the cellular phenotypes they represent. Finally, it discusses system design and implementation based on expert feedback. Two real-world use-cases demonstrate the applicability of our approach to cutting-edge cancer research.

## 2 Related Work

### Cellular Screening and Phenotype Analysis

A wide variety of biomedical visual analysis (VA) tools have been developed for analyzing images of cells grown in culture. CellProfiler [14] is a widely-used open-source tool for cell-based screening that can segment and quantify cells grown in multi-well dishes using preconfigured pipelines. Related open-source and commercial tools include, Phaedra [53], HCS-analyzer [50], ScreenIt [22], Cytosplore [33], PerkElmer’s High Content Profiler [24], and Genedata Screener [27]. Few of these tools are applicable to tissue imaging where image size is typically very large, the number of different cell types is substantial, and cells are crowded closely together in a complex environment. Tissue images require a richer interactive and EDA framework in which experts can bring in domain knowledge to steer automated algorithms and evaluate results. The existing tool closest in performance to Facetto is HistoCat [60], a Matlab-based toolbox with a visual front-end for analyzing tissue images acquired using imaging mass spectrometry. The developers of HistoCat are part of our team and were involved in the Facetto evaluation. Facetto extends concepts developed in HistoCat by providing a web-based interface, by integrating machine learning (ML) into the data analysis workflow and by adding multiple forms of data faceting.

### Multi-modal Analysis of Image and Feature Data

Several VA tools explicitly focus on image data and feature data in combination. Bannach et al. [6] use image data to extract radiometric features from radiology data that can be used for grouping patients into cohorts. Corvlo et al. [18] extract features from medical image data for digital pathology, but the tool is not configured to support multi-channel images. Other multi-modal analysis approaches include tumor tissue analysis [57], medical perfusion data [28, 49] and time-varying volumes [26]. However, none of these tools are designed to work with large, high-resolution multi-channel tissues images and they do not support spatial selection and interactive ML-based analysis. Facetto also has new tools for faceting data and for visual guidance (via phenotypic trees) in the form of orienting, as defined by Ceneda et al. [15].

### Interactive Clustering and Classification

Clustering, an unsupervised ML technique, enables the discovery of new patterns in data, based on a defined distance function and cluster strategy [37]. Supervised classification techniques learn dependencies from examples and apply the knowledge to categorize new data [23]. Both approaches have been combined with interactive interfaces to select input subsets [46], actively steer algorithms [32, 34], understand model internals [63] and input-output relations [66], and to explore, compare [41, 42] and refine results [17]. Users are frequently involved in these processes to add domain knowledge and make final decisions for critical applications.

A less explored aspect of ML is combining unsupervised and supervised learning methods, that are “[…] two complementary concepts embedded at the very heart of visual analytics” [64]. Combining them can be especially advantageous when little or no ground truth is known and there is a repetitive need to classify new data similarly. An example of a simple combined approach is classifying new instances according to nearest cluster centers (nearest neighbors) [7, 31]. A more advanced combined application involves radial basis function (RBF) networks in which cluster centroids determine the centers of input or middle-layer neurons [16, 35]. Other approaches [19, 48] leverage clustering for selecting representative subsets in active learning, taking into account natural breaks in the data distribution. Thom et al. [10] provide both clustering to find spatio-temporal patterns and topic-based classification in a VA setup but it applies them in isolation. With Facetto we go one step further by iteratively and interactively combining clustering and classification to support reasoning [70] (Section 6.4).

### Hierarchical Data Faceting and Provenance

A variety of methods exist to visually capture data provenance and track consecutive filtering actions during a data manipulation and analysis process. Many of these approaches follow a filter-flow metaphor [69]. Prominent analysis tools include KNIME [7] and Orange [21]. SOMFlow [59] can iteratively facet and refine clusters, with the history of previous decisions visualized in a flow graph. Alternatively, DataMeadow [25] subdivides (facets) datasets into nodes, each representing a data subset whose instances and features can be visually encoded within the node. VisTrails [12] provides visualizations to create and edit dataflows and support provenance management. Following the provenance types defined by Ragan et al. [56], Facetto supports recall and action recovery, enabling data subsetting and facetting and also storing the cognitive outcome and information derived from the analysis process. Scatterblogs [10] also supports data subsetting, using classifiers, for social media analysis. However, unlike Facetto, it does not enable learning from subsets asnd application of the knowledge so gained to create new sets in the hierarchy.

### Large-scale Image Viewing

Many medical and biological visualization systems focus on the display of 2D imaging data. Deep Cell Zoom [3] and the Cancer Digital Slide Archive [13] support online browsing of curated biological datasets. However, both tools are pure image viewers and do not enable interactive manipulation of viewing parameters other than zooming and panning. To make image viewing and rendering scalable to large data, multi-resolution techniques such as image pyramids [30] must be employed. Facetto is based on OpenSeadragon [51], a scalable web-based framework for viewing multi-resolution images. We additionally support high-precision (32-bit) segmentation masks, multi-channel rendering, and multiple rendering modes (Section 5.2). Multi-channel rendering is conceptually similar to multi-volume rendering [11, 67], in which multiple volumes are blended together to form a final output image, for example, using one dataset for defining opacity and another dataset for defining the output RGB color [44]. Facetto supports similar rendering modes, by blending multiple channels or using a single channel (e.g., a segmentation mask) to define the rendering mode for the other channels.

## 3 Multiplex Tissue Imaging Workflow

Figure 2 shows an overview of the CyCIF imaging process.

**Fig. 2.**
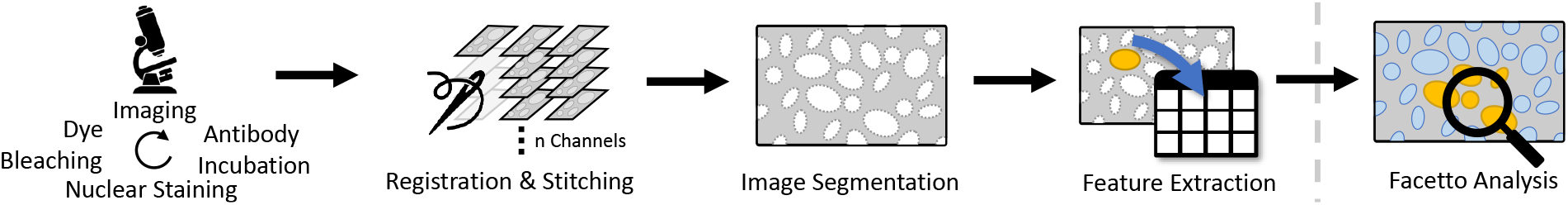
The CyCIF imaging workflow. Steps 1-4: Acquisition and Preprocessing. Tissue sections are stained with antibodies chemically linked to a fluorophore; typically 3-5 antibodies are combined. The sections are imaged and fluorophores are then chemically inactivated (bleached) making it possible to perform another round of staining and imaging. Multiple cycles allow collecting images with 60 or more different ‘‘channels,” each of which represents the staining pattern from a single antibody. Images are collected in successive tiles, which are then stitched together. Cells are segmented, and features such as total intensity per cell are extracted. The resulting data are analyzed interactively in Facetto (step 5).

### Acquisition

Multiplex tissue imaging uses tissue samples recovery from patients for diagnosis (biopsies) or during surgery (resections). Following sectioning (typically into 5 micron thick slices) samples vary in size from a few square mm to several square cm. Almost all approaches to multiplex tissue imaging use antibodies against specific proteins to identify cells types, visualize structures within these cells, and measure the levels of specific proteins involved in regulation of cell state (the targets of such antibodies are often referred to as “markers”). In the specific case of CyCIF, chemically fixed tissue samples are stained with a mixture of antibodies each of which is chemically linked to a different fluorescent molecule (a fluorophore). Tissue samples with bound antibodies are then imaged using a high-resolution optical microscope, resulting in 16-bit four-channel images that show precisely where each antibody has bound, and thus, the locations and levels of proteins of interest. Finally, fluorophores are chemically inactivated (bleached) and another round of incubation with fluorescently-labelled antibodies is performed (often called “staining”) followed again by imaging. This process is repeated multiple times (see Fig. 2, step 1), leading to 60 or more images of the same tissue.

### Processing

High resolution optical microscopes have limited fields of view. Thus, larger samples are acquired as a series of image tiles, which are then stitched together to form a complete image (see Fig. 2, step 2). Different fluorescence channels are registered to each other using software such as the open-source tool ASHLAR [5]. This yields precisely aligned images, 30,000 pixels in each dimension, that are then segmented to identify individual cells (see Fig. 2, step 3). Segmentation is a challenging task, currently performed using a random forest classifier [62], but is subject to continuous improvement. Segmentation information is stored in 32-bit masks that define the cell ID for each pixel in a multi-channel image stack. Next, per-cell mean intensities, area, shape, and neighborhood features are extracted for 10^6^ or more individual cells per specimen (see Fig. 2, step 4). The resulting multichannel images, segmentation, and high-dimensional feature data are then ready for interactive analysis in Facetto (see Fig. 2, step 5).

### Terminology and Data Characteristics

For the remainder of this paper, we assume that a CyCIF dataset contains (1) a multi-channel tissue image, (2) a segmentation mask, and (3) a feature table of image-based features in csv format. Each *image* (or *channel*) in the multi-channel image stack represents data from a different antibody stain. The segmentation mask spatially locates individual *cells* in each tissue specimen and assigns cell *IDs*. Each row in the csv file (i.e., *instance*) is identified by its cell ID and also contains the extracted feature values for that cell. *Features* represent either expression levels (i.e., average intensity for specific cell in a specific channel), subcellular morphological features, or spatial features involving multiple cells.

## 4 Goal and Task Analysis

To better understand the goals and tasks associated with the analysis of multiplex tissue (CyCIF) data, we conducted in-depth, semi-structured interviews with ten domain experts. All of them are affiliated with the Harvard Laboratory of Systems Pharmacology. Two experts are oncologists (hereinafter referred to as O1, O2), two are pathologists (P1, P2), and six are cell and computational biologists (CB1-6). Additionally, we participated in weekly group meetings over several months during the development of Facetto and the acquisition of initial datasets.

### 4.1 User Roles and Goals

**Oncologists** are physicians who diagnose and treat cancer and have direct contact with the patients they treat. The oncologists involved in this project are also research scientists active in understanding the molecular basis of diseases such as melanoma, breast, and lung cancer. Their primary clinical goal is to identify optimal treatment regimens for individual patients; their primary research goal is understanding the molecular basis of drug response and resistance.

Pathologists are physicians who analyze tissues and biological samples to diagnose disease. Anatomic pathologists primarily use microscopes in their work and have unique expertise in understanding the complex features of histological preparations that are diagnostic of specific diseases. The pathologists involved in the project are also research scientists and, within the CyCIF team, have a special role in guiding and checking work performed by other investigators and by computer algorithms. Their primary clinical goals are distinguishing diseases from each other (differential diagnosis) and working closely with oncologists on treatment strategies; their primary research goal is understanding the role played by tumor cell type and state and the tumor microenvironment in disease processes and response to therapy. **Cell and Computational Biologists** combine biomedical knowledge with skills in technical subjects, ranging from mathematics and physics to computer science. Within this project they are responsible for developing the CyCIF methods, for collecting primary data and for processing this data to create stitched images, segmentation masks and sets of derived features (Section 3). They develop analysis scripts and apply a variety of computational methods to image and numerical data. The cell and computational biologists involved in this project are trained in interpreting the morphologies of cells, but they do not have the pathologists’ deep knowledge of tissue structure and disease.

The three user roles are dependent on each other, as cell and computational biologists are typically guided in their interpretation of tissue data by physicians and physicians are dependent on biologists for wet and dry method development, data transformation, and analysis. Facetto is designed to support close collaboration and interaction among pathologists, oncologists, and cell and computational biologists.

### 4.2 Tasks and Challenges

From interviews with experts, we derived a series of tasks that need to be performed on images to meet the overall goals of understanding cancer biology and diagnosis of disease in individual patients. Along with these tasks, we identified gaps and challenges that currently hinder the analysis of tissue imaging data.

#### T1: Cell Type Discovery and Calling

The task most frequently mentioned by all experts is identifying and analyzing specific types and states of cells based on the intensity and pattern of staining with specific antibodies (O1, O2, P1, P2, CB1-6). *Challenges:* Challenges lie in processing, displaying, and faceting the large and high-dimensional data, as well as in mutual support of manual and automated analysis. A lack of adequate tools makes this task very time-consuming at present.

#### T2: Overview-Detail Exploration of Multi-Channel Image Data

A crucial task for oncologists and pathologists is rapid navigation and visualization of multi-channel images (O1, O2, P1, P2). Pathologists are accustomed to moving slides back and forth physically on a microscope stage and switching between high and low power views. They rely on a seamless visual experience to make a diagnosis. *Challenges:* Image analysis must not only support seamless pan and zoom, but also switching between groups of channels. Current tools do not scale beyond 4-5 channels and lack on-demand rendering, blending of channels, and means to emphasize (and recall) regions or individual cells of interest.

#### T3: Data Filtering and (Sub-)Structuring

Another task frequently performed by pathologists is *gating*, which refers to manual filtering of selected image channels based on the channel’s intensity value range (often visualized as a frequency-intensity plot), or specific spatial features or regions of interest (P2, O2, CB2). *Challenges:* Analysis steps such as gating are often applied in an iterative manner in which the data is hierarchically faceted into subsets. These subsets can then be further analyzed, used in benchmarks, or exported for presentation or reuse with other samples. Thus, tracking the evolution and provenance of gates and gated data is important.

#### T4: Proofreading and Analyzing Results in Spatial Context

Many algorithms operate on features computed from images following segmentation; these include mean intensity value per cell and channel. Feature extraction and segmentation from tissues, in which cells of different sizes and shapes are crowded together, are challenging tasks for which software tools are still being developed. As a result, it is essential that the results of feature extraction are checked and corrected prior to downstream data processing (CB1, CB3). This requires effective means to link feature and image space (P1, O1). *Challenges:* Currently, such linking is only supported by HistoCat [60], and generally requires domain experts to continuously switch between tools (CB2).

#### T5: Deriving Profiles for (Sub)regions and Classes

Once a type/region is detected, it is important to identify, annotate, and extract a profile of typical marker distributions within an area of interest (O2, P2). The profile includes statistical measures and distributions of cell features and can be used to present the outcome of an analysis session, diagnosis, or as a starting point for further analysis. *Challenges:* The variables used to construct profiles, and the ways in which these variables are displayed, are not standardized and can only be developed by human-machine interaction.

## 5 Visual Exploration of CyCIF Data

We used the tasks of Section 4 to guide the design and implementation of Facetto, playing the *translator* role put forth in the design study methodology by Sedlmair et al. [61]. Figure 1 gives an overview of Facetto’s interface. Using the image viewer, users can explore spatial features (**T2**), hierarchically facet the data into subsets (**T3**), and look at profiles of cell phenotypes and spatial regions (**T5**). Classification and clustering can be triggered in different views to support the hierarchical discovery of cell types (**T1**). To proofread and analyze results (**T4**), we have implemented features such as image tool-tips and an interactive sortable table for cell features. The table view gives details on individual cells and supports manual manipulation of cell feature data. A key design goal is preventing feature overload and effectively guiding users through specific, repetitive tasks. The Facetto image viewer is therefore surrounded by tools that can be activated and deactivated as needed. We now present and justify these design and implementation decisions in more detail.

### 5.1 Data Faceting and Hierarchical Analysis

Cell type discovery and labeling (**T1**) is an iterative process. Experts usually start with a region of interest (ROI) at high resolution (so that individuals cells are visible) that represents a subset of the complete multi-channel image stack and then define spatial and image features to create data subsets of interest (**T3**). The results obtained on this ROI are then applied to the entire specimen. We support multiple iterations of this process with hierarchical data faceting. Users can build up a hierarchy of different data subsets, and we automatically display this ongoing analysis in a hierarchical phenotype tree view. This allows users to track their progress, and to maintain an overview of their data faceting and analysis steps they have performed (**T5**) (i.e., allowing users to look up results from past analysis steps to recover or re-execute them with a different image or in a different setting).

Figure 3 shows the visual representation of our faceting approach, a so-called *phenotype tree*, an interactive n-tree visualization. At the beginning of an analysis session, the tree holds a single root node representing the entire dataset. From region and feature-based selections, users are able to create new subsets and store them as child nodes in the tree. We have integrated algorithms (Section 6) that make it possible to further divide the active data subset into classes and clusters, which can again be stored as new nodes in the tree. In contrast to many automatic data lineage systems, we allow domain experts to manually decide which subsets are meaningful and to add them to the phenotype tree. We provide a context menu that makes it possible to activate a node’s subset (i) for use as the current data (sub)set across all views in Facetto (ii) to delete a subset/node, and (iii) to create new nodes, e.g., from selections or classification results. To improve semantic understanding and recall, users can name, color, and annotate each selection. Facetto can also derive visual profiles (**T5**) for each discovered (sub)region and class. Figure 3 shows an entire phenotype tree (left) and an exemplary snippet (right) with an active node (denoted by the presence of a halo) containing active selections (orange fill line showing the number of cells in the selection). It descends from a spatial selection and is the parent to a clustering result with three detected subgroups, representing different immune cell types. All views in Facetto support this faceting approach by being able to display either the entire dataset, the active subset, or the current selections a user is working with.

**Fig. 3.**
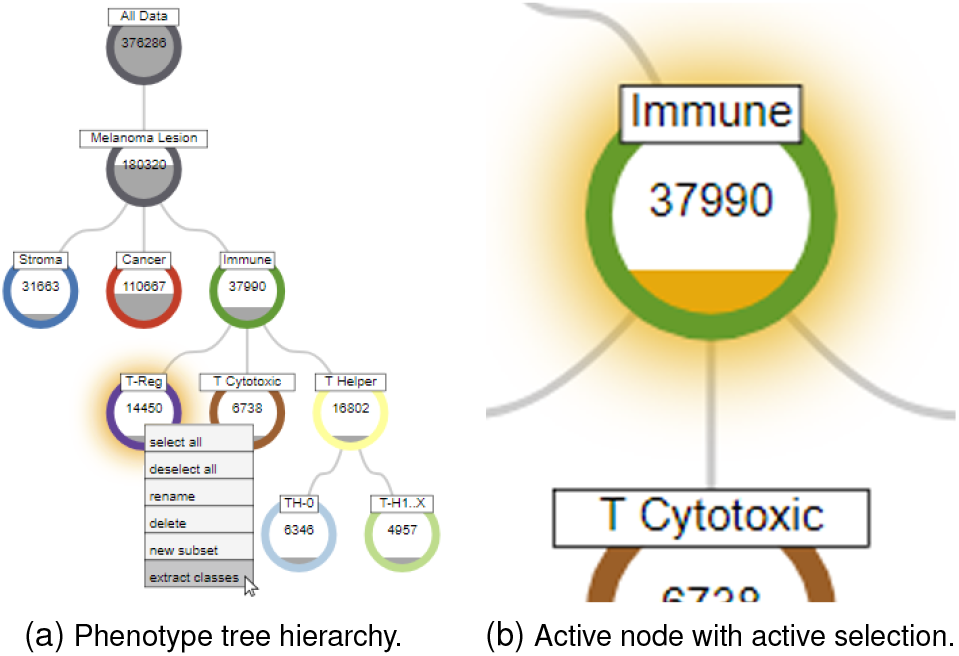
The phenotype data flow tree supports the analyst in the phenotype analysis workflow. Users make selections (e.g., by filtering, clustering, or classification) in an initial dataset (i.e., root). Selections can be stored as subsets and applied in a hierarchical manner, leading to a tree structure that maintains analytical provenance. Each node has a label, color, and displays the number of contained cells as a fill line.

### 5.2 Image Exploration

Facetto’s image viewer allows users to navigate and explore large multichannel image data (T2) using a scalable multi-resolution visualization approach. The visualization supports interactive and seamless zooming and panning, on-demand rendering of multi-channel information, manual selection of cells, and highlighting of classification and clustering results as well as faceting operations. The image viewer also provides details on demand when hovering over individual cells.

#### Image Data

Each image tile in CyCIF is a 16 bit grayscale image, typically comprising 4 × 10^6^ pixels (the dimensions of a scientific grade CMOS camera). Each channel is recorded in a separate grayscale image that is registered to other channels and pseudocolored for visualization. Segmentation assigns an ID (cell ID) to each cell in the stitched image. We store segmentation masks at the same resolution as the image data but in a 32-bit integer format to support the large number of cells in each image. A resection specimen imaged in 60 channels with its associated segmentation masks represents 60 GB of data.

To support interactive image exploration we employ a multiresolution rendering approach. In a pre-processing step, an image pyramid is computed for each channel by repeatedly down-filtering the original data. We use bicubic downsampling to maintain smooth gradients in the lower resolution images. We also create an image pyramid of the cell segmentation mask. To prevent incorrect interpolation of cell IDs at lower image resolutions, we perform downsampling by nearest neighbor filtering. To ensure fast data loading times, each resolution level in the image pyramid is further split into smaller image tiles of a fixed resolution (e.g., 130 × 130 pixels) and stored on disk. Figure 4 shows our different image pyramids and rendering options.

**Fig. 4.**
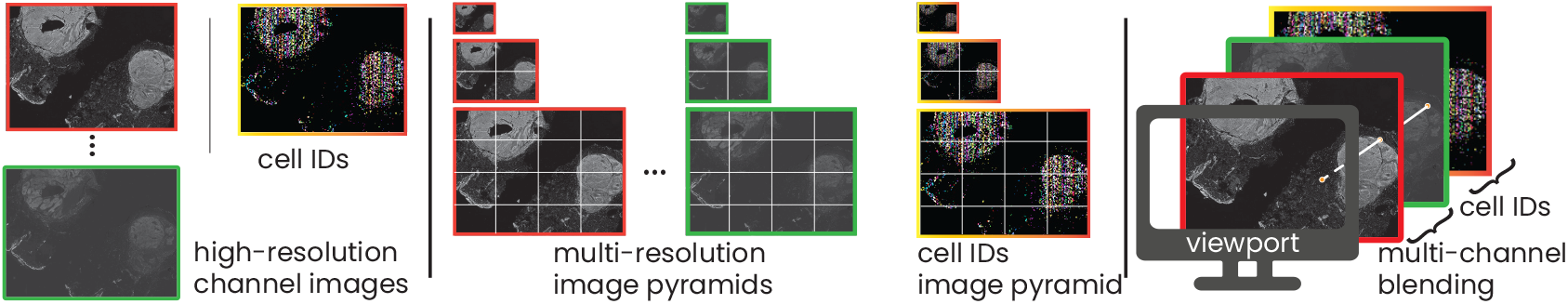
Multi-resolution image rendering. Left: The input to our rendering pipeline is a stack of high-resolution images (one image per CyCIF channel) and a segmentation mask containing a cell ID per pixel. Middle: We pre-compute multi-resolution image pyramids for all input data. Right: During rendering, we blend channels and choose between different rendering modes based on a pixel’s cell ID, e.g., for highlighting a selected set of IDs.

#### Multi-Resolution Image Viewer

Our image viewer is based on OpenSeaDragon [51], an open-source, web-based image viewer library. To support the visualization of CyCIF data, we added support for image tile caching, segmentation masks, multi-channel rendering, and advanced selection and filtering capabilities. During rendering, the image viewer determines the required tiles for the current viewport, zoom level, and active image channels. If the tiles are not in the cache (i.e., they have not been loaded yet, or have been purged), the tiles are requested, loaded, and transferred to the viewer. Tiles are rendered asynchronously, which ensures that the application remains responsive even if very large datasets are being loaded in the background or if the network connection is slow. The visualization is updated automatically whenever a new tile has been loaded. We also support interactive transfer functions (i.e., color look-up tables) to allow users to adjust the visual representation of the displayed image data. This is likely to become increasingly important as color mapping of histological data is subject to government regulation [52]. Figure 5 shows a single-channel image (Fig. 5a) as well as different zoom levels in a multi-channel rendering (Fig. 5b-d) of a CyCIF dataset with 376,286 cells.

**Fig. 5.**
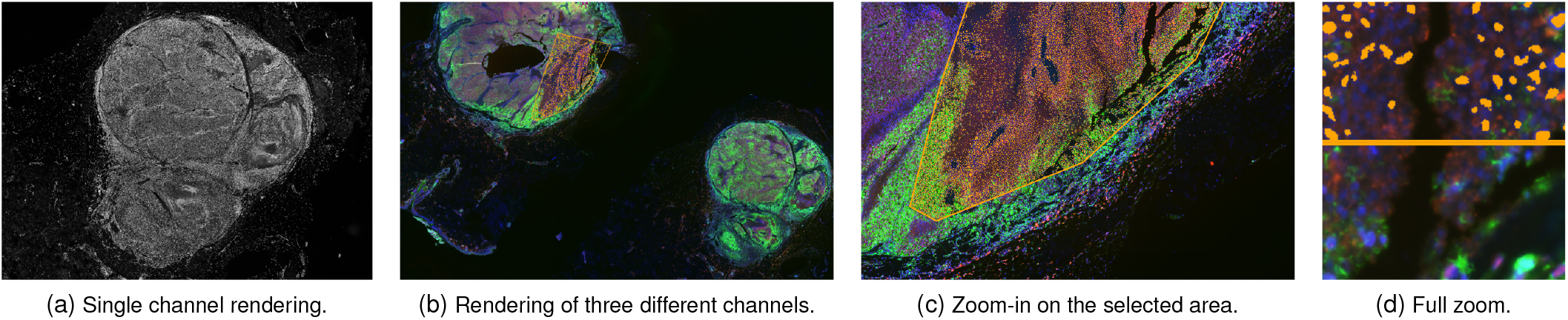
Image viewer render modes. (a) shows single channel rendering, while (b-d) show different zoom levels of a 49.8 GB dataset. The rendering shows three channels (proteins *pS6, HES1* and DNA) with a selection on a metabolically active area. (d) Top: Selected nuclei are highlighted in orange. Bottom: 3-channel rendering without nuclei highlights.

#### Multi-Channel Rendering

To analyze the image data (**T3, T4**), we visually encode features of interest and subsets of the data. To make spatial distributions and correlations among multiple channels pre-attentively visible, we allow the user to blend the data from different channels into a single image (see Fig. 5b-d). Users select the appropriate channels for their current tasks based on domain knowledge. For multi-channel rendering, we first retrieve pixel intensities for each (x,y) position in the selected image channels and then apply linear color and opacity transfer functions to the individual channels. We blend the contributions of the individual channels to get the final RGBA (i.e., color plus alpha) output for that pixel. Facetto supports different blend modes of the individual channels, such as alpha compositing, addition, or multiplication [55]. When mapping up to three concurrent channels (default colors are red, green, and blue) Facetto can guarantee nonoverlapping colors. However, after feedback from domain scientists, we added support for up to five concurrent channels, which allows users to look at additional channels that have no spatial overlap. For each channel, users can set and modify a linear color and opacity transfer function by specifying the respective intensity range (i.e., lower and upper bound) as well as the colors of the transfer function.

A caveat of multi-channel rendering is that mixing different colors at pixel level makes detailed interpretation of the resulting pixel color difficult, as RGB colors are not distinct perceptual channels [47, 65]. However, after speaking to our experts we found that this multi-channel color mapping is state-of-the-art in cell-based image analysis, and our domain scientists expect, and heavily use, this feature in visualization tools. Furthermore, domain experts most commonly combine channels that have almost no spatial overlap (e.g., two antibodies that stain different types of cells or non-overlapping structures). Usually, three of the five possible mixed channels are well separated visually, and five channels are still informative if spatial overlap is limited. To assist in evaluating overlap, users can quickly toggle the visibility of individual channels. Figure 5b-d shows a multi-channel visualization of channels *pS6* (a ribosomal protein whose phosphorylation state is a measure of cell signaling), *HES1* (a transcription factor) and *DNA*.

#### Cell-based Rendering

The tasks of interactive exploration and filtering (**T2, T3**) also require support for dynamic selection of cells and their visual display in image space. This requires a means to convey that cells in a ROI are part of a currently selected subset. To support the visual selection and highlighting of individual cells, we use the segmentation mask (Section 3) to determine the cell ID associated with each pixel. Next, we dynamically adjust the render mode based on the current pixel’s cell ID. Facetto supports different render modes, depending on whether the user is currently focusing on the entire dataset, a subset (i.e., a node in the hierarchical phenotype tree), or a user selection (i.e., based either on manual selection or a clustering/classification result). Facetto can then display individual cells in their original grayscale intensity, apply a color and opacity transfer function (see Fig. 5d, bottom), or show color overlays for cluster/class membership (see Fig. 5d, top).

#### Focus and Context

To reveal contextual feature information for an individual cell or for a selection, users can click on a cell in the image viewer and show a visual profile card in which data statistics are summarized (**T5**). The card shows a boxplot of the feature space, the phenotype labels, and a short summary that includes any previous user annotation, making it possible for information to be acquired sequentially over a number of sessions involving multiple users.

### 5.3 Feature Space Exploration

The image viewer makes it possible to blend different image channels to reveal spatial structures and patterns, but analyzing the multivariate dependencies in the data remains difficult. During the image preprocessing step we extract features for each segmented cell (Section 3) such as mean intensity for each cell in each channel, cell area, shape, and neighborhood features. To enable in-depth analysis of these features (**T3, T4, T5**), we provide a set of visualizations that display the feature values and value ranges for an active selection of cells. We also represent cell similarity and distributions for subsets and selections in feature space. Finally, we provide means to investigate and manipulate individual cell values in an interactive tabular display.

#### Exploring Feature Distributions

For the exploration of each channel’s distribution, we integrated a ridgeplot (see Fig. 6) that comprises multiple area charts alongside relevant information about the variable being examined (**T5**). Each chart represents a feature’s value distribution (a ridge). Typically, the distribution of features that are derived from a channel’s intensity values is skewed, having a few distinct peaks and some outliers. We, therefore, apply square root scaling to make dense regions in the intensity distribution more visible and reduce visual peaks. Each ridge is equipped with range sliders, allowing to filter the underlying data with visual feedback at the level of the distribution (**T3**). The selection range is used directly for specifying the color transfer function applied in the image viewer. To allow the exploration of feature distributions for subsets of the image or individually selected cells, we overlay the parallel x-axes in the ridgeplot with vertical polylines (see Fig. 6, orange paths), encoding each cell’s features as a connected path, similar to parallel coordinate plots [36]. To reduce clutter, we decrease opacity as the number of selected cells grows, an approach that provides an indication of the correlations and distribution ranges of a selection, while focusing less on individual cells. Alternatively, a single polyline can be used to represent the mean values of a cluster.

**Fig. 6.**
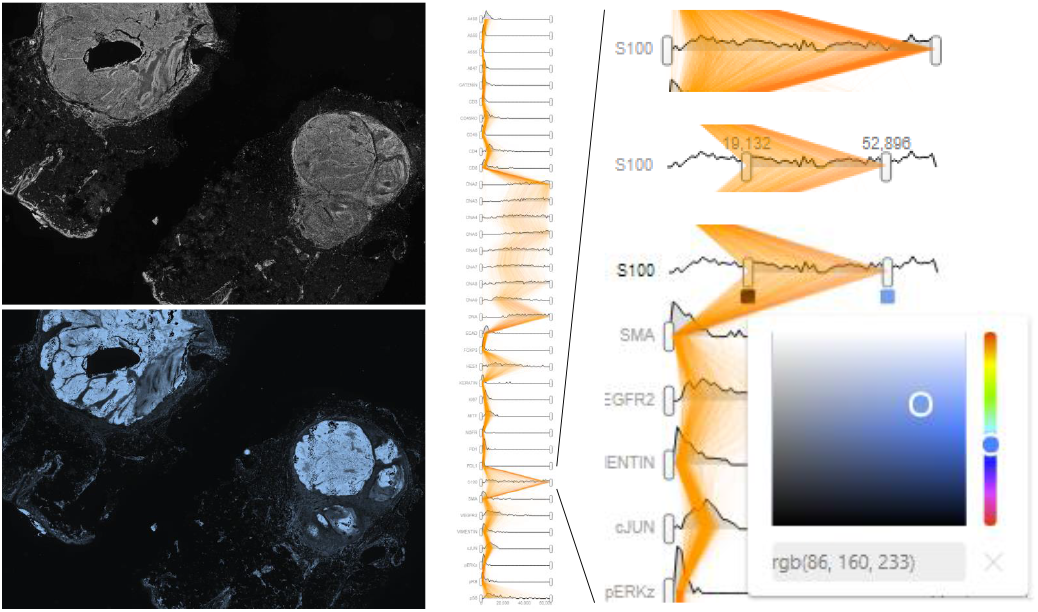
The ridgeplot (middle and right) shows intensity distributions for each of the 44 image channels, arranged on parallel x-axes. Feature values for the currently selected cells are visualized using vertical polyline overlays (in orange). Users can interactively filter individual channels, select channels for display in the image viewer (left), and adjust transfer functions used for rendering (bottom left and right).

#### Exploring Individual Cell Features

Experts can sort, inspect, select, and manipulate individual values for each feature of a cell using the interactive visual tabular display (see Fig. 1f). The main goal of the tabular view is a) detailed analysis and direct manipulation of individual cells, and b) allowing users access to the original data table. We encode the extracted intensity values in the tabular view as numbers, as well as by using small multiples of bars. The tabular view is also color encoded, with each color indicating a distinct phenotypic class. All views in Facetto are connected via brushing and linking so that users can analyze a selection from different perspectives. In this way, users can mark (and edit) individual cells with certain features in the tabular view and inspect spatial context in the image view or vice versa (**T4**).

#### Dimensionality Reduction of High-Dimensional Features

We visualize higher-order similarities and differences between data subsets (**T5**) using dimensionality reduction techniques and subsequent display in a 2D scatterplot. We use UMAP (uniform manifold approximation and projection for dimension reduction) [45], a recently developed machine learning technique, to display features of the current data subset (see Fig. 7b). This algorithm is similar to the popular t-SNE projection [43] but preserves more of the global structure and has superior run time performance [68]. We have found that UMAP works particularly well on multiplex-immunofluorescence data from tissue sections [68]. Users can also switch to displaying the first two components of a PCA (principal component analysis); this is faster to compute for very large datasets. The data is visualized in gray, and active selections are shown in orange or are visually encoded by their phenotype (class/cluster) color. The projection reveals how well certain cell groupings cluster or separate from each other. Our domain experts are very familiar with dimensionality reduction techniques. Hence, this is a valuable means to discover and review phenotypes and to discover novel cell sub-populations with similar staining patterns. We support filtering of cell groupings in our projection views by polygon selection, similar to the image-based selection mechanisms (Section 5.2).

**Fig. 7.**
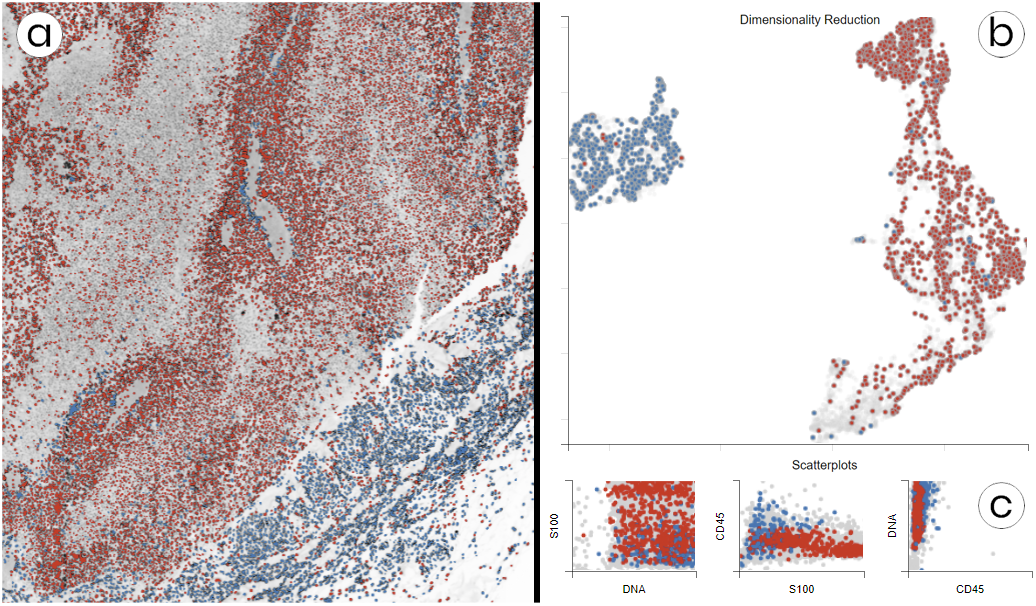
(a) Image viewer showing a region with colored cell nuclei from clustering, overlaid onto the DNA channel. (b) UMAP projection scatter-plot of an active subset with color-coded clusters. The clustering separates cancer regions (red) from immune cells (blue). (c) Small-multiple scatterplots show correlations of active rendering channels (here *DNA*, the tumor marker *S100*, and immune cell marker *CD45*).

## 6 Semi-automatic Phenotype Analysis

We developed a novel semi-automatic approach to cell state analysis for CyCIF data. It supports data faceting (**T3**) with unsupervised and supervised machine learning methods that are tightly integrated into Facetto’s visual interface. This allows domain experts to steer, review, and manipulate the outcome of those automatic methods (**T4**). Specifically, we leverage clustering as a means to discover novel cell subtypes (**T1**) and acquire new knowledge, and classification as a means to propagate the learned cell types/states across CyCIF images (**T1**). We close the loop by allowing users to set the output of one method as the input to the other method in an interactive manner.

### 6.1 Unsupervised and Supervised Learning Loop

Both clustering (unsupervised) and classification (supervised) facet data into subsets by assigning labels to cells. These labels denote *categories or classes* and are completely independent of the cell IDs used by the image segmentation mask (Section 3). When there are many unknown aspects of an image and dataset (e.g., a different patient, a new type of cancer, or new immuno-fluorescence antibodies), a bottom-up analysis strategy [54] is often considered the best approach. Clustering is then a helpful means to identify similar cell types and states and to segregate them from each other. After carefully reviewing and adjusting the clusters for a smaller subset, a user can then apply the optimal *faceting strategy* to other cell subsets. This is accomplished by using the labeled data to train a CNN model and subsequently apply it to new data. Having a dataset classified and faceted into broad categories (e.g., cancer, immune, and stromal cells), users may want to further subdivide the data to arrive at more fine-grained subtypes. This may include subdividing all immune cells into their different myeloid and lymphoid subclasses by examining antibody staining of cell lineage markers (e.g., *CD3* for T cells, *B220* for B cells, etc.). Clustering in Facetto runs on feature data, while classification runs on the original image data. This makes the classification step more expensive, but has the advantage of incorporating data from full high-resolution images.

To limit cognitive load in the approach outlined above, Facetto allows users to track and review machine learning results and decisions and quickly access subsets of the data on which to train and apply ML methods by means of hierarchical faceting and the phenotype tree.

### 6.2 Expectation Maximization Clustering

To assist exploratory discovery of cell types, Facetto includes EM clustering [20], a partitioning method that iteratively computes maximum likelihood estimates using a Gaussian mixture model. The method assumes that the data is sampled from an underlying statistical population that can be expressed by an analytic distribution. Each iteration consists of an expectation step *E* and a maximization step *M. E* estimates the distribution of the underlying population from an available sample, given a certain fixed model. *M* finds means and variances of Gaussian mixture components that maximize expected log-likelihood of the observed data and hidden variables. These steps are alternated until the log-likelihood changes fall below a given threshold. EM clustering has several advantages over other clustering approaches, such as k-means and DBSCAN: It can find clusters of different size, density, and shape. However, Facetto also allows easy integration of other methods.

Figure 8b depicts our clustering pipeline with each cell represented by features. A user first defines the input instances (cells) and features to be considered. We normalize the selected features using a *log*_10_ transform and percentile normalization, which are standard in singlecell image analysis. In percentile normalization, we transform [0.1%, 99.9%] values to [0, 1], and truncate outliers. We then apply EM clustering with *k* clusters, where *k* is selected by the user. The results can then be reviewed and manipulated. Users can apply domain knowledge to re-label certain cells or to further split clusters into smaller subsets (subtypes). To judge cluster quality, users can inspect cluster separation in high-dimensional space using UMAP (see Fig. 7b) and investigate silhouette coefficients across multiple clustering runs.

**Fig. 8.**
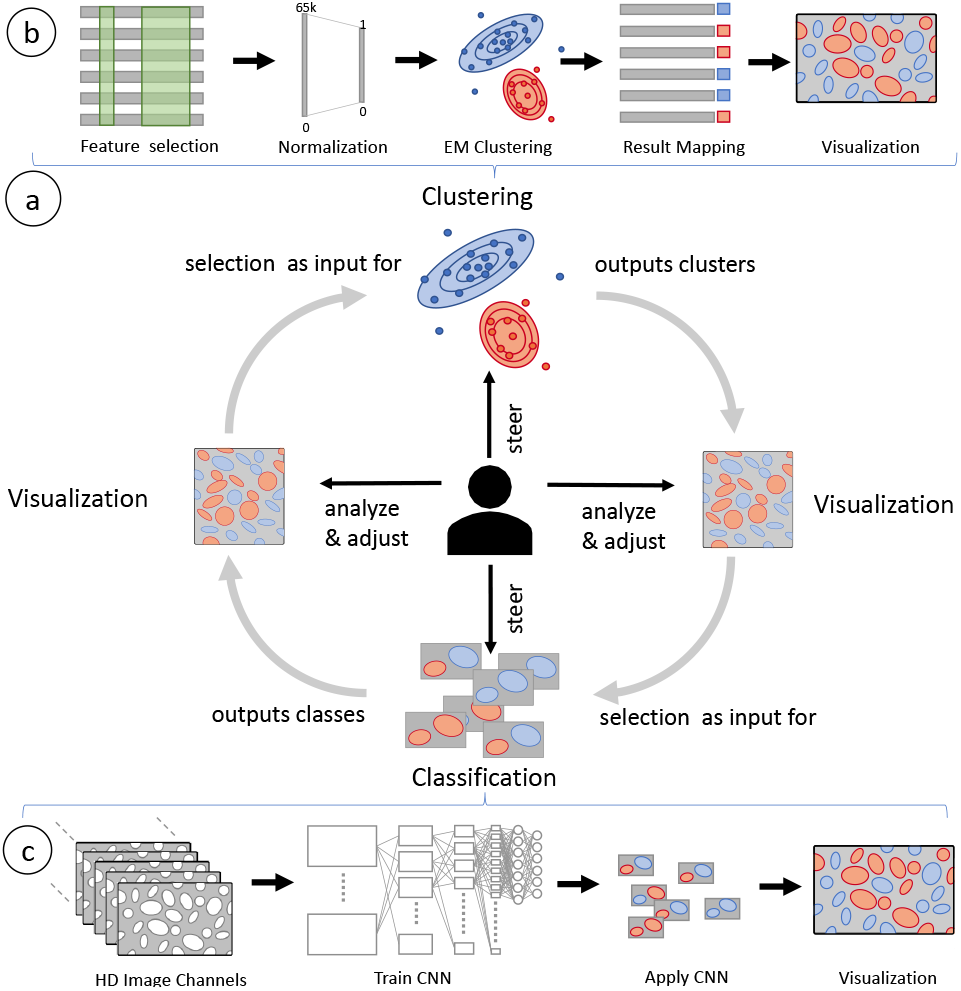
(a) Facetto allows users to employ clustering and classification in an iterative (loop) setting where users can use the result of one to steer the other, and vice versa. (b) EM Clustering: User-selected data is normalized, clustered, and finally visualized. (c) Classification: Multichannel images of labeled cells are used as input to train a CNN. For selected cells, respective image tiles are inputs for the classification. Classified cells are extracted and directly visualized in the image viewer.

### 6.3 Classification via Convolutional Neural Network

To classify cells based on pre-defined types as well as those discovered through clustering, we have integrated a deep-learning convolutional neural network (CNN) into Facetto. Figure 8c illustrates our cell classification pipeline. The inputs are a multi-channel input image and a segmentation mask of cell IDs. The network yields probability maps for each cell type as an output. We convert the probability maps into classification labels for each cell by applying a softmax function and aggregating cell-type probabilities for each segmented cell.

#### Network architecture

Our CNN consists of an encoder and a decoder with skip connections [58] to exploit low-level and high-level features. The encoder has 8 convolution layers with 3 × 3 filters and 3 pooling layers, decreasing spatial resolutions to increase receptive fields. We built the decoder using 9 convolution layers, 9 filters, and 3 deconvolution layers. The deconvolution layers restore the original spatial resolution that had been decreased by the pooling layers.

#### Training

We train the network by accepting images of cells and their ground truth cell type labels as annotated by experts, such as ‘cancer,’ ‘immune,’ and ‘stroma.’ This ground truth can be derived from manual labeling or from the result of a previous EM clustering run. Due to computational complexity, we divide the entire image into tiles and load only the tiles with ground truth data to train the network. We minimize a softmax cross entropy between predictions and their ground-truths using the Adam optimizer [38]. Parameters in the network are randomly initialized and the learning rate is set to 0.001. To judge the quality and trustworthiness of the trained classifier we display the prediction accuracy and precision and recall in a confusion matrix.

#### Application

Once trained, users can apply the classifier to selected data subsets. As in the training step, a user defines a set of cells to be classified. We then compute which tiles need to be loaded (i.e., all tiles that contain any of the selected cells), using a quad-tree acceleration structure. We then classify each cell in the loaded tiles. Subsequently, the results are filtered to match only the requested labels to be displayed. Users can visually inspect results and manually sort out or relabel questionable cell classifications.

### 6.4 Expert in the Loop

Facetto is an “expert in the loop” system designed to exploit the domain knowledge of pathologists and cell biologists while automating routine tasks and supporting quantitative analysis that is not possible by eye alone. In Facetto, experts configure automated approaches (e.g., by selecting input instances and features) and directly analyze results. Algorithms can be triggered by active selection through the visual interface. When the result is available, the interface updates to show results and also provide contextual scales and information on accuracy. By default, each new cluster or class is assigned a name (label) and a distinct color from a categorical color scale. This color can be used to display cluster membership of cells in the image viewer. Alternatively, users can toggle the render mode to multi-channel rendering of the original image channels. Users can move back and forth between an image viewer providing information on the spatial distribution of the cell types (classes/clusters) and various feature-based visualizations revealing the distribution of image-derived information in high-dimensional space. For example, if UMAP indicates that clusters 1 and 2 are well separated, while cluster 3 is more connected to cluster 1, users can rerun the clustering with a different number *k* of clusters. Brushing and linking allows users to drill down to individual instances, and to review and relabel selected values and assigned classes in the image and table view.

## 7 Implementation

Facetto is a web-based client-server system. The back-end is implemented with Flask, a python microframework for web development. The stateless back-end provides a restful interface, making it possible to retrieve image and feature data, and steer analytics. Facetto’s components are based on a modular design, where new computational methods can be plugged in, and called by defining new restful endpoints. Clustering, classification and UMAP computation are implemented in Tensorflow [1]. Clustering, training, and classification of 10^3^ cells takes 0.05, 25, and 20 seconds respectively. We intend to further optimize training time to enable a more interactive experience. Facetto’s front-end runs in the web browser and is based on Javascript, HTML, CSS and D3.js. The image viewer extends OpenSeaDragon (see Section 5.2) for client-side dynamic rendering in the browser. Scalable scatterplots are implemented in hardware-accelerated HTML canvas views.

## 8 Evaluation

We present two case studies that demonstrate how Facetto can accelerate and improve the analysis of multiplex tissue images. Both evaluations were carried out in a small meeting room with Facetto displayed on a wall monitor. We collected think-aloud feedback and logged user analysis steps and actions. We also collected regular feedback on the tool’s usability from a computational biologist using Facetto for her research. In the case studies described below, we followed the visual analytics evaluation protocol developed by Arias-Hernandez et al. [4].: while the domain experts guided and steered the analysis, we operated the user interface. This approach allowed us to evaluate the capabilities and limitations of our approach on cutting edge research tasks and questions rather than simply collect feedback on details of the interface.

### 8.1 Use Case 1 - Exploratory Data Analysis (EDA)

In this use case, two pathologists (P1, P2) and one cell and computational biologist (CB3) used Facetto to analyze samples of lung tumor over a span of ninety minutes. The main goal of our experts was to freely explore the data and thereby gain novel insight into the tumor biology by leveraging linked image and feature representations. The users were also interested in evaluating the accuracy of cell segmentation.

#### Dataset

We analyzed a dataset of lung tissue, containing a stack of 44 images (channels), a nuclei segmentation mask, and a table with extracted mean intensity values per cell for each channel. Each channel had a dimension of 14,448 × 11,101 pixels, resulting in 13.6 GB raw channel data, and contained 110,500 successfully segmented cells.

#### Proofreading

The team started the analysis by looking at the segmented cells in the image viewer. We activated the channel corresponding to nuclear DNA and then added other channels. Our users had not previously been able to interactively explore segmentation masks aligned with image data. Using Facetto, users were able to identify immediately some segmentation errors involving incorrect cell merging and splitting (**T4**). Using the tabular data view, they sorted the cells based on their size, used brushing to select the largest cells, and then inspected these cells in the image viewer. These “large” cells were commonly ones in which the segmentation algorithm had failed to separate individual cells that were in close contact. Users then utilized clustering (based on cell size) to analyze different types of segmentation problems (i.e., boundary errors vs. overlapping nuclei). Using Facetto, it was simple to exclude these cells from subsequent analysis.

#### Phenotype Analysis

Next, our users explored simple cell phenotypes within the lung specimens. To spot tumor and immune cells (**T1**), they leveraged Facetto’s multi-channel rendering. Users first looked at the distribution of nuclei (bright structures in the *DAPI* channel), and a single tumor marker (*Keratin*). Next, they added the channel for a protein that marks proliferating cells (*Ki-67*) and for a stromal cell marker (*aSMA*). By toggling channels on and off, it was possible to confirm the expected patterns of co-staining and marker exclusion.

Next, one expert identified an area with a high concentration of immune cells lying at the border of the tumor (**T2**). By activating different channels related to immune cell markers (*CD4, CD8a, FOXP3*), users were able to visually assess the spatial distribution of two functionally distinct types of immune cells (Cytotoxic T-cells and Regulatory T cells). The ratio of these cell types is widely thought to play a role in the susceptibility of different cancers to the latest generation of immuno-oncology drugs, and is therefore important to estimate reliably. Our users then selected a spatial region in the heart of the tumor with the polygon tool and turned to clustering. They asked us to run clustering on markers for tumor cells (*Keratin*), immune cells (*CD45*), and cell size, to distinguish tumor and non-tumor cells (**T3, T4**). We clustered the tumor subset further based on *Ki67* staining and iteratively increased the cluster number to *k* = 5. After confirming in the image viewer that these five clusters corresponded to different types of proliferating tumor cells (as judged by toggling color-coded cluster overlays), we labeled corresponding nodes in the phenotype tree and then studied the nodes in more detail (**T5**). By looking at the UMAP, users confirmed a clear separation among clusters.

#### Feedback

Users were excited about Facetto’s proof-reading and analysis capabilities and found the interface more intuitive and responsive than any existing tool. They stated that the interaction between image viewer and feature analysis was essential for identifying meaningful combinations of overlapping cells. The pathologists were more skeptical than other users about automated classification and its applicability to infer cancer subtypes, but considered it a useful way to examine large datasets. The ability to interactively review classification results is likely to be important for these users. Users relied heavily on Facetto’s multi-channel rendering and observed that the flexible transfer functions involving use of a histogram to set upper and lower bounds in the rendered image helped a lot (CB3). The color overlay of cluster results was also perceived as intuitive, since Facetto uses visually distinct colors for transfer functions and clustering. To reduce clutter in ridgeplots, users wanted to toggle between a detailed view (one vertical polyline per cell) and an overview (one polyline per cluster).

### 8.2 Use Case 2 - Ovarian Cancer Phenotype Analysis

In a second use case we analyzed images of ovarian cancers over a span of two hours, guided by a gynecologist/oncologist (O2), and a medical informatician with six years of experience in cancer research (CB4). The goal was to explore ovarian cancer-specific spatial patterns.

#### Dataset

Our users examined two datasets involving 40-channel images of serous ovarian cancer resections. The first dataset (D1) contained 5,273 successfully segmented cells and was 5,666 × 9,306 pixels in size (4.1 GB). The second dataset (D2) contained 358,380 segmented cells and was 22,703 × 14,841 pixels in size (26.4 GB).

#### Proofreading

After visual exploration of dataset D1 at different zoom levels in the image viewer (**T2**), we marked a specific spatial region from which to retrieve feature details in the tabular view. To the experts’ surprise, when we highlighted the resulting cell IDs from the ROI query in the image view, the cell IDs did not align with the selected region (see Fig. 10). Upon closer inspection, we identified an error in the extracted feature data (**T4**). Evidently some cell IDs were dropped during preprocessing; this arose because virtually all of the software tools used for tissue imaging are in active development. Spotting an error like this is crucial; when the extracted cell features do not match the cell IDs in the segmentation mask, subsequent analyses are incorrect. CB4 stated that without Facetto, they might not have discovered the error. Our users then switched to dataset D2 with no preprocessing errors.

#### Phenotype Analysis

The users explored the image view and chose channels that would highlight cell nuclei (*DNA*) and a marker of proliferation (*Ki67*) that is high in rapidly dividing cancer and immune cells. Our collaborators discovered areas of the tumor with different numbers of proliferating cells (**T2, T3**). To analyze a specific region of the sample in greater detail, users selected a ROI using the polygon selection tool, and then stored the resulting set of cells as a node in the phenotype tree (see Fig. 9b, 9f). To distinguish between cancer and non-cancer cells, they applied integrated EM clustering on *DNA* and *Ki67*, with *k* set to two clusters (**T1**). Facetto displays the clustering results visually as colored overlays on the cells in the image viewer (see Fig. 9c). Hiding the cluster overlays allowed experts to fine-tune the color transfer functions for individual channels (**T3**). User O2 further refined the clustering by adding *E-Cadherin*, a marker responsive to tumor cells. Subsequently, CB4 stated that the clusters look a lot more accurate than those identified by other approaches (see Fig. 9g) and she verified that the UMAP displayed expected features.

**Fig. 9.**
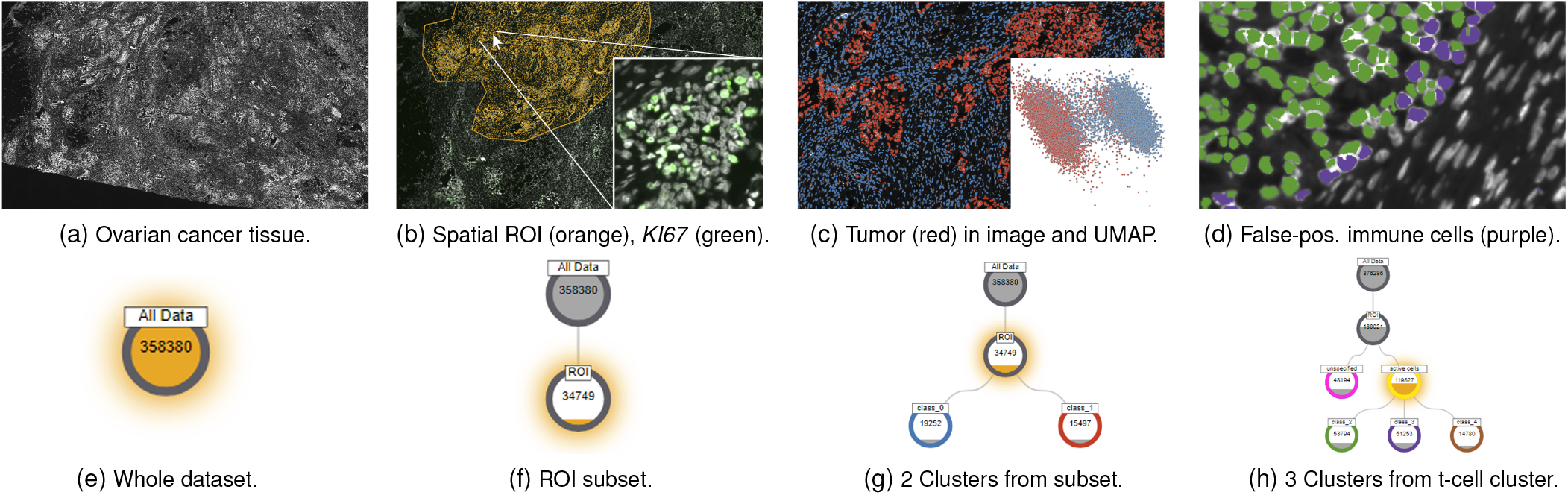
Ovarian cancer phenotype analysis (use case 2). Left to right: Experts start with a high-level view of the data (a, e), extract a ROI (b, f), cluster cells in the region into cancer and immune subpopulations (c, g), and subsequently subdivide one of the clusters into subtypes (d, h).

**Fig. 10.**
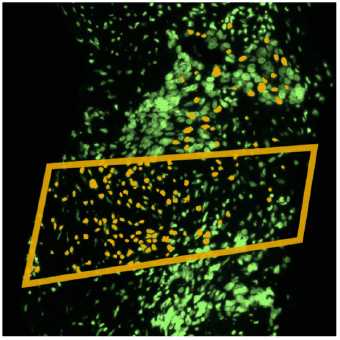
Mismatched cell IDs in dataset D1.

Next, our users attempted to identify different cell sub-phenotypes by analyzing staining patterns in different image channels (**T1, T3**). We started a new phenotype tree, extracted dividing cells into a cluster, and then created further subsets from that cluster node using the hierarchical faceting view. Our experts clustered on *DNA, PAX8, CD4, CD3, CD8a, CD163, PD1, IBA1, CK7, CD11b, E-Cadherin*, and *Vimentin*. By looking at *CD3* in combination with the colored cluster overlays, users were able to identify putative false positives in the *CD3* staining as well as T-cells of particular interest (see Fig. 9d, 9h). They then added textual labels to the clusters in the phenotype tree. Next, they wanted to use the cell clusters they had discovered as the basis for further analysis of the rest of the image. This included classification, exporting findings, and performing a more in-depth statistical analysis (**T5**).

To showcase Facetto’s proofreading and classification capabilities, we introduced the experts to the editing mode in the tabular data view as a means to validate individual cell-cluster memberships. We also demonstrated how cells can be manually reassigned to a different cluster (**T4**). This data can then be used to feed the updated cluster and phenotype labels as ground truth data into a classifier (Section 6.4). Our experts found this especially promising, considering the large number of samples they currently need to label and proofread manually.

#### Feedback

O2 and CB4 were excited about the integration of visualization and ML. They stated that this integration sped up the workflow significantly. It allowed them to identify different cell types, and to fine-tune clustering by adjusting the ROI and selected image features. Importantly, they could also directly check for segmentation errors and mapping issues between image and extracted feature data, which would be hard to detect otherwise. A key advantage of Facetto over other tools for these applications is that it is possible to concurrently review both, original image data and the results of image analysis. Users found the phenotype tree view to be a particularly useful means for exploring identified subsets and hierarchically analyzing data.

### 8.3 Lessons Learned and Limitations

#### Tool complexity

One general drawback of VA tools used for multidimensional image data is their inherent visual complexity. Our collaborators repeatedly requested more functionality, even if that came at a higher cognitive load. We accommodated this by allowing users to toggle between visualizations involving different levels of complexity. For example, color encoding in the image view is usually based on transfer functions for selected channels, but can also be used to show cluster/class membership using visually distinct colors. Other examples of variable complexity included toggling the vertical polylines in the ridge view from showing one line per cell to one line per cluster.

#### Accommodating different users

Facetto supports users with different medical and technical backgrounds. The Facettto UI launches into a common ground view comprising the image viewer and channel selection in the ridgeplot. Additional tools, such as clustering or hierarchical phenotype analysis, are arranged around the image viewer, and can be used depending on the user’s background and analysis goals. All of our current users are highly involved in research, but in a production environment Facetto could support multiple roles, such as histotechnician and physician, each with access to different functionality.

#### Building trust

Trustworthiness is an important aspect to data visualization. Our collaborators always want access to the original data. Facetto supports this in the tabular view and in the image viewer. Furthermore, we allow scientists to examine results of ‘black-box’ ML algorithms, crucial for building trust. One current limitation of Facetto in this regard is that our phenotype tree does not support fuzzy sets to represent partial memberships to classes (e.g., 80% probability of cancer). Additional trust could be built with visual guidance pointing users to interesting channels. In practice, however, current users often know which channels to examine, based on prior knowledge.

## 9 Conclusion and Future Work

We describe a newly developed visual analytics tool, Facetto, that combines unsupervised and supervised learning for hierarchical phenotypic analysis of multi-channel tissue images. Facetto addresses an acute need for software that complements manual analysis of histological data, which is the norm in a clinical setting, with automated analysis. Facetto as an open-source tool allows others to add algorithms and interfaces as needed. With Facetto, we hope to substantially improve our ability to interpret complex tissue images in both research and translational settings, thereby advancing our understanding of disease and of new and existing therapies. In the longer term we expect tools like Facetto to be applied in a clinical setting as a means to improve patient diagnosis and enable personalized therapy.

User testing suggests that Facetto is already superior to many existing image analysis tools. In the future, we intend to add features that allow users to evaluate and improve other aspects of the tissue imaging workflow, including verification and correction of segmentation results with active learning. Ultimately, Facetto is expected to support ML methods that transcend the limitations of conventional segmentation methods. We also intend to add new means for creating and applying classifiers and for leveraging GPUs to parallelize computation for capturing user feedback in real-time.

Of particular importance in tissue imaging is exploiting the ability of anatomic pathologists to identify image features that have scientific and diagnostic significance. By interactively linking numerical and spatial features derived from images, machine learning and image visualization, Facetto provides a “human-in-the-loop” approach to accelerating and improving image exploration and analysis. With Facetto, we aim to let physicians, oncologists and cell and computational biologists benefit from each other’s expertise: physicians can leverage computers to cope with data overload and computational biologists can identify and test algorithms for automating repetitive tasks. Ultimately, we expect that the vast majority of work in analyzing multiplexed image data will be automated, allowing scientists to focus on interpretation and innovation. A critical aspect of this transformation will be making the results of ML interpretable to users and also subject to user-guided improvement. This type of interactivity is at the heart of the Facetto approach.

## Acknowledgments

We thank Rumana Rashid, Shannon Coy, Sandro Santagata, Anniina Farkkila, and Julia Casado for their valuable input and feedback. We thank Benjamin Izar, Jia-Ren Lin and YuAn Chen for data and advice on the manuscript, and Denis Schapiro, Clarence Yapp, Jeremy Muhlich for image stitching and segmentation. This work is supported by the Ludwig Center at Harvard Medical School, by NCI Grant U54-CA225088, by King Abdullah University of Science and Technology (KAUST) and the KAUST Office of Sponsored Research (OSR) award OSR-2015-CCF-2533-0.

